# Single-cell foundation models predict durable CAR T response despite imperfect cell annotation

**DOI:** 10.64898/2026.07.22.740224

**Authors:** K. Leo Shen, Zhiliang Bai, Mingyu Yang, Na Li, Rong Fan

## Abstract

CD19-targeted chimeric antigen receptor (CAR) T cell therapy achieves high initial response rates in B-cell acute lymphoblastic leukemia (B-ALL), yet half of patients relapse within one year. Pre-infusion product composition decoded by single-cell RNA sequencing (scRNA-seq) carries information predictive of long-term CAR T persistence, but extracting this information from individual patients typically requires highly sophisticated bioinformatics expert annotation, limiting clinical translation. Here, we evaluate whether single-cell foundation models (scFMs) can extract clinically actionable information from engineered CAR T products. We applied four scFMs (scGPT, scFoundation, CellPLM and UCE), including fine-tuned versions of scGPT and scFoundation, to paired basal and CD19-stimulated pre-infusion CAR T products from 33 pediatric patients with B-ALL. Although annotation accuracy declined relative to healthy peripheral blood references, scFM-derived cell composition stratified patients with long-duration B-cell aplasia with a leave-one-out cross-validated area under the receiver operating characteristic curve of 0.879 (95% confidence interval, 0.742–0.986). Notably, foundation-model-identified cell proportion analysis matched or exceeded expert annotations for several predictive features, demonstrating that accurate clinical prediction may not require perfect per-cell annotation to begin with. CD8+XCL1/2+ cells were further identified as the biomarker consistently associated with durable CAR T persistence across models under CD19 stimulation, whereas other candidate populations showed limited reproducibility. Finally, we translate these findings into a locally deployable decision-support AI agent that predicts the probability of sustained CAR T persistence from pre-infusion CAR T scRNA-seq data.

## Introduction

Chimeric antigen receptor (CAR) T cell therapy targeting CD19 has transformed the treatment of relapsed or refractory B-cell acute lymphoblastic leukemia (B-ALL), with complete remission rates exceeding 70% in pediatric patients^1–3^. Despite this efficacy, approximately half of treated patients relapse within the first year, and the molecular determinants of durable response have been reported^4–6^ but highly variable and difficult to pinpoint on the individual patient basis, in particular, prior to the administration of cellular immunotherapy. B-cell aplasia (BCA), the sustained depletion of normal B cells in peripheral blood, serves as a clinically validated pharmacodynamic surrogate for CAR T cell persistence, with patients exhibiting early B-cell recovery within six months almost invariably experiencing relapse^7^. A small subset of patients sustains BCA for years, and recent work has demonstrated that elevated type-2 functionality in the pre-infusion CAR T product is significantly associated with long-duration BCA exceeding eight years^8^. These observations argue that the pre-infusion product itself encodes signatures predictive of long-term clinical outcome, yet extracting these signatures at single-cell resolution typically requires bespoke bioinformatics expert annotation that limit reproducibility and clinical translation.

Several reasons explain why the pre-infusion product encodes outcome. CAR T cells are manufactured from autologous patient T cells through *ex vivo* activation, viral transduction, and expansion, and the resulting product carries the imprint of the patient’s prior antigen exposure history, T cell differentiation state, and exhaustion burden. Products enriched for less-differentiated T cell subsets (naïve and central memory) tend to expand more robustly *in vivo* and persist longer, while products dominated by terminally differentiated effector cells or by exhaustion-marker-positive subsets tend to lose function early^9–12^. The ratio of CD4 to CD8 cells, the abundance of regulatory T cells, and the proportion of cells in active proliferation at the time of infusion are all candidate determinants of post-infusion behavior. Profiling these compositional features at single-cell resolution before infusion therefore offers a direct window onto the biological substrate of CAR T persistence but realizing that promise depends on profiling methods that scale beyond bespoke expert annotation.

Single-cell foundation models (scFMs), which are large transformer-based neural networks pretrained on tens of millions of cells, have emerged as a candidate solution to this annotation bottleneck^13–16^. By learning a generic representation of cellular state from broad transcriptomic atlases, models such as scGPT^13^, scFoundation^14^, CellPLM^15^ and UCE^16^ promise to enable cell type identification, reference mapping and downstream phenotype prediction without dataset-specific retraining. Reported applications span batch integration, perturbation prediction and zero-shot annotation across diverse tissues^17^. Notable additional scFM architectures not evaluated here include Geneformer^18^ and scBERT^19^. Since pretraining at this scale captures a generalizable representation of cell state that applies across datasets and architectures, it is often assumed that biological inference from scFM embeddings should be similar regardless of model choice.

However, whether this promise extends to clinically realistic settings is contested. Several recent benchmarking studies have reported that current scFMs do not reliably outperform classical methods on standard tasks, especially in zero-shot inference and under distribution shift^20–22^. The published literature has focused on aggregate annotation accuracy on healthy reference atlases, leaving critical questions for translational application unaddressed. First, do scFMs retain useful resolution on engineered T cell products that have been *ex vivo* manufactured and antigen-stimulated—biological contexts substantially distinct from the resting peripheral blood compositions of pretraining corpora? Do annotation imperfections negatively impact downstream clinical prediction, or can scFM-derived cell type proportions support outcome stratification despite imperfect labeling? Lastly, do different scFM architectures, evaluated on the same dataset, identify the same biomarkers, in a reproducible manner for deployment?

Here, we show that scFMs can extract clinically meaningful signal from pre-infusion CAR T products despite imperfect per-cell annotation. We evaluated four scFMs (scGPT, scFoundation, CellPLM and UCE) in zero-shot mode and two of them (scGPT and scFoundation) after fine-tuning, using a single-cell atlas of pre-infusion CD19-CAR T products from 33 pediatric B-ALL patients enrolled across the first two CAR T trials at the Children’s Hospital of Philadelphia (NCT01626495 and NCT02906371)^8^. The cohort is stratified into three BCA outcome groups by duration of post-infusion B-cell aplasia: BCA1 (short, n=17), BCA2 (intermediate, n=11), and BCA-L (long-duration sustained aplasia, n=5). Each product is profiled under both basal and CD19-stimulated conditions, and performance is contextualized against a healthy peripheral blood mononuclear cell (PBMC) reference atlas^23^ and a post-infusion *in vivo* dataset from a single patient^24^ scFM zero-shot accuracy declines by 0.21–0.25 weighted F1 units when moving from PBMC to the engineered CAR T product, with antigen stimulation imposing an additional systematic penalty. Despite imperfect per-cell annotation, model-derived proportions discriminate BCA-L from non-sustained BCA groups with leave-one-out cross-validated area under the receiver operating characteristic curve (AUC) of 0.879 and, for multiple specific cell types, outperform ground-truth-derived proportions in discriminative power. CD8+XCL1/2+ cells are the only feature reaching permutation significance in both scGPT and scFoundation under CD19 stimulation, identifying a candidate biomarker for sustained CAR T persistence. Cross-model agreement on biomarker identity is partial, and several clinically important populations, including CD4+ regulatory T cells and post-infusion exhausted CD8+ subsets, are unable to show statistically significant association across all four architectures. We package these findings into a locally deployable clinical decision-support agent that returns a per-patient predicted probability of long-duration BCA.

## Results

### Single-cell foundation models resolve immune taxonomy in a healthy PBMC reference atlas

To systematically evaluate the translational potential of scFMs for CAR T cell therapy, we established a unified benchmarking pipeline encompassing gene vocabulary harmonization, model-specific preprocessing, embedding generation under zero-shot or fine-tuned settings, and reference-based cell type annotation using FAISS-accelerated k-nearest-neighbor (k-NN) mapping in the learned embedding space (Fig. 1a). We evaluated four representative scRNA-seq foundation models spanning distinct pretraining strategies and architectures: scGPT, a generative transformer pretrained on approximately 33 million human cells; scFoundation, a masked autoencoder trained on over 50 million transcriptomes; CellPLM, a cell language model pretrained on both single-cell and spatial transcriptomic datasets; and UCE, which leverages protein language model-derived gene representations to generate universal cell embeddings. Because these models differ substantially in training objectives, model architecture and pretraining data, cross-model comparison provides an opportunity to evaluate the robustness and reproducibility of biological inference across current generations of scFMs.

**Fig. 1:**
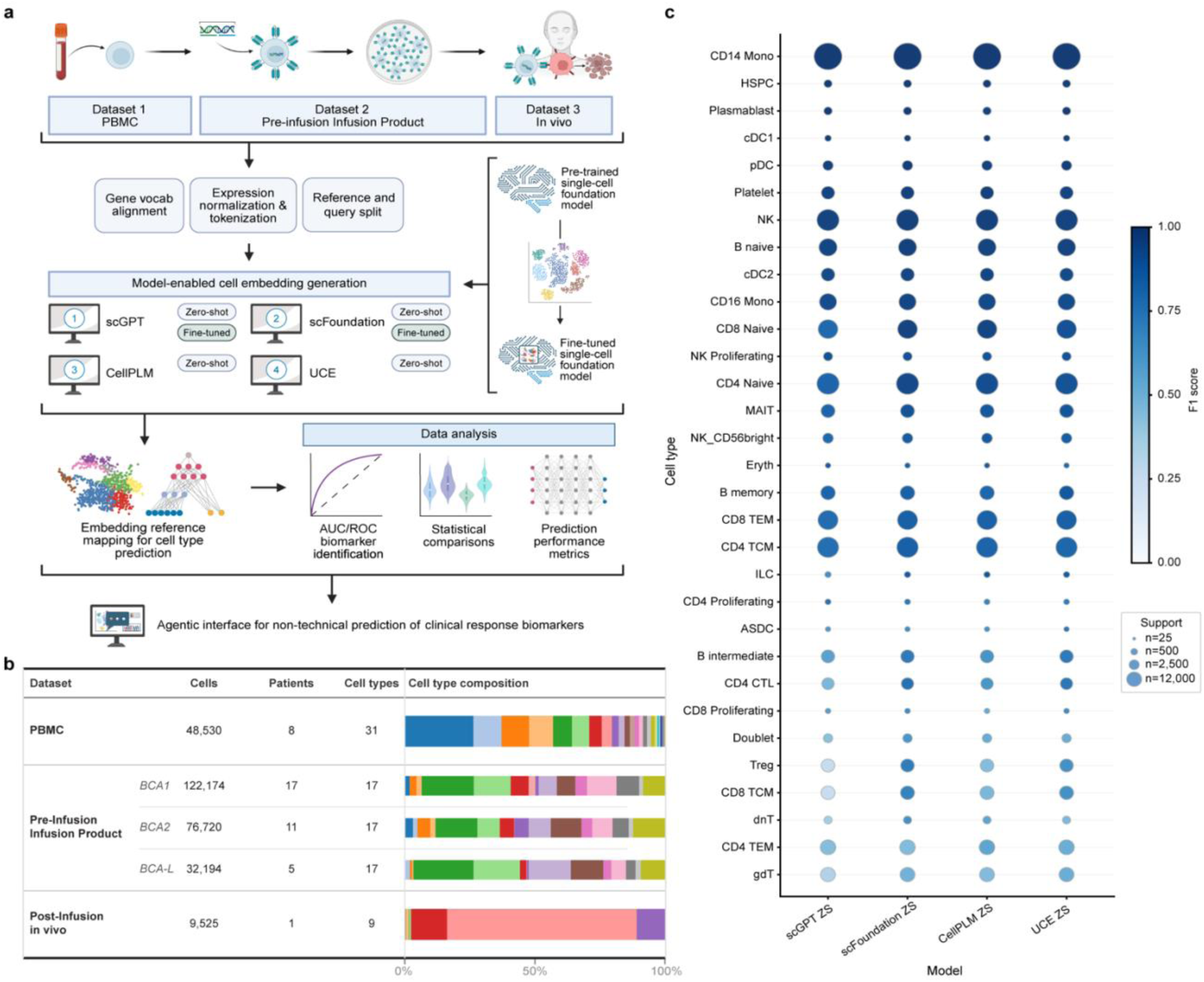
Evaluation framework and PBMC reference performance of single-cell foundation models. a, Schematic of the evaluation pipeline. **b,** Overview of the three evaluation datasets, including total cell counts, patient counts, number of annotated cell types, and cell type composition. **c,** Per cell-type zero-shot F1 scores across all four models on the PBMC dataset. Created in BioRender. Shen, L. (2026) https://BioRender.com/vu0p0nq is licensed under CC BY 4.0.

Three complementary datasets were selected to progressively evaluate foundation model performance across increasing levels of biological and clinical complexity (Fig. 1b). We first used a healthy PBMC reference^23^ atlas comprising 48,530 cells from eight healthy donors with 31 expertly annotated immune cell populations, providing a well-established benchmark for immune cell annotation. We next analyzed a clinically annotated pre-infusion CAR T cell product cohort^8^ consisting of 231,088 cells from 33 pediatric patients with relapsed or refractory B-ALL enrolled in the first two CD19 CAR T clinical trials at the Children’s Hospital of Philadelphia. This cohort is uniquely valuable because each infusion product was profiled under both basal and CD19-stimulated conditions, while long-term clinical follow-up stratified patients into three clinical response groups according to the duration of post-infusion BCA, a validated surrogate of CAR T persistence. Finally, we evaluated a post-infusion peripheral blood dataset^24^ comprising 9,525 CAR T cells collected *in vivo* following CAR T therapy to assess model generalizability beyond the manufactured infusion product into the dynamic post-treatment state. Together, these datasets enabled evaluation from healthy immune cells to engineered cell therapy products, and ultimately to CAR T cells functioning *in vivo*.

We first established the upper bound of model performance using the healthy PBMC reference. All four foundation models were evaluated in zero-shot mode and achieved weighted F1 scores exceeding 0.84, indicating that pretrained embeddings accurately capture canonical immune cell identities without task - specific retraining (Fig. 1c). Highly abundant immune populations, including CD14⁺ monocytes, naïve B cells, natural killer cells and naïve CD4⁺ T cells, were annotated with near-ceiling accuracy across all four architectures. In contrast, rare immune populations, including γδ T cells, double-negative T cells, innate lymphoid cells and CD4⁺ effector memory T cells, exhibited substantially lower performance regardless of model architecture. Annotation accuracy therefore closely tracked cell abundance, indicating that current scFMs remain constrained by the same class-imbalance limitations observed in conventional reference-mapping approaches rather than overcoming them through large-scale pretraining. Notably, this reference atlas is a widely used public dataset and may be represented in the pretraining corpora of one or more of these models; if so, the near-ceiling performance on abundant populations represents a best case, and the persistent difficulty with rare populations despite that potential exposure further indicates that pretraining scale does not resolve class-imbalance limitations. This healthy PBMC benchmark establishes a reference ceiling against which performance on engineered CAR T products and subsequent translational analyses can be interpreted.

### CAR T product composition imposes a systematic accuracy penalty for foundation models

We next asked whether foundation models pretrained largely on unmodified immune cell atlases remain effective when applied to engineered CAR T infusion products, a substantially different biological context. Unlike circulating PBMCs, CAR T products undergo *ex vivo* activation, viral transduction, expansion, and manufacturing before infusion. Consequently, their learned embeddings fail to capture the unique signaling programs induced by synthetic CAR receptors, which differ fundamentally from native T-cell receptor signaling in amplitude, timing, and downstream effector coupling^25, 26^. The CAR T dataset therefore represents a clinically relevant domain-shift benchmark that more closely reflects the conditions under which scFMs would ultimately be deployed. Across all four models, zero-shot annotation performance decreased substantially relative to the healthy PBMC reference, with weighted F1 scores declining by 0.21–0.25 across all BCA outcome groups (Fig. 2a). This consistent reduction across architectures indicates that the engineered transcriptional state of CAR T cells, rather than any individual model design, presents a fundamental challenge for current foundation models.

**Fig. 2:**
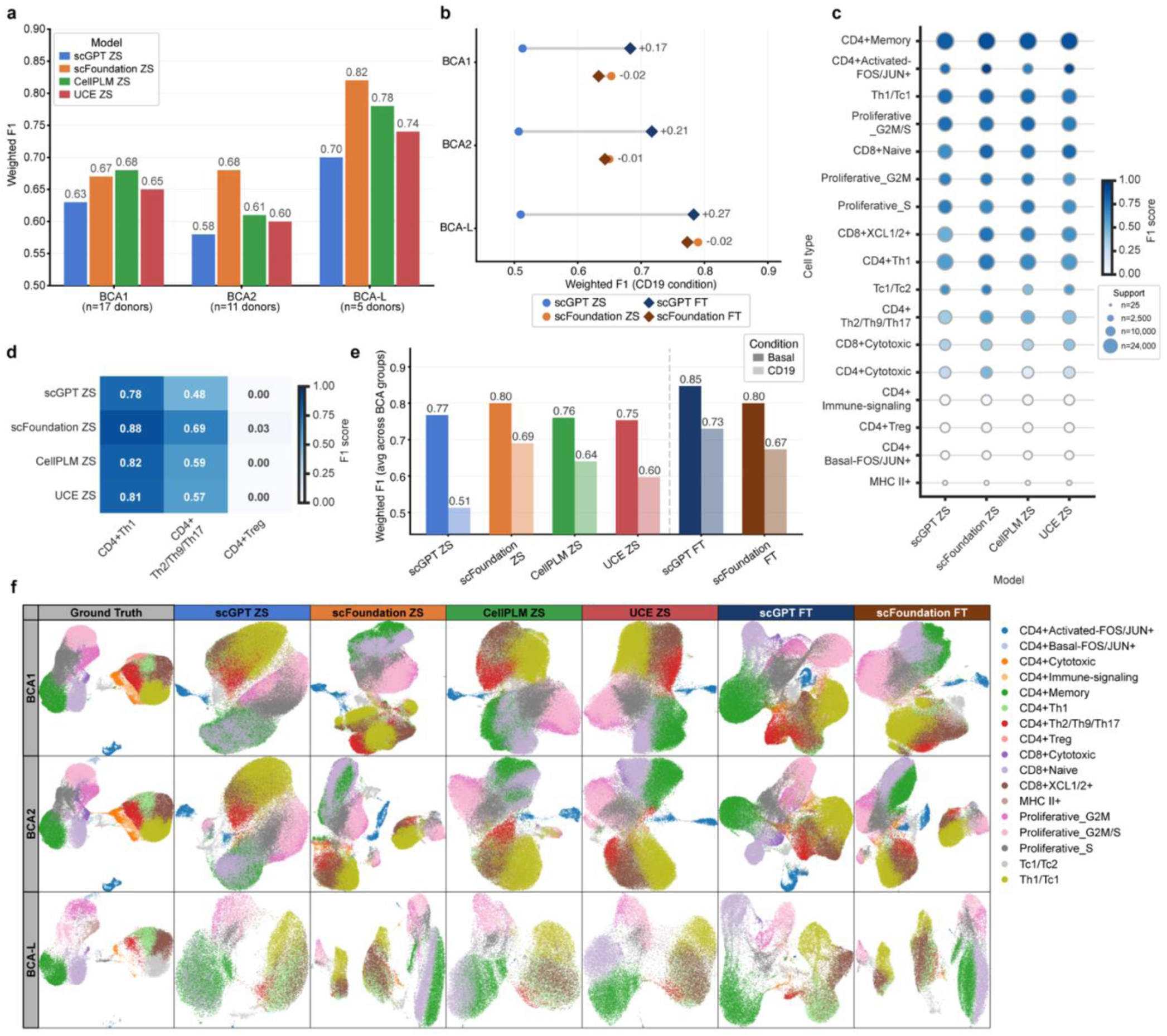
Zero-shot annotation performance on the pre-infusion CAR T infusion product. a, Weighted F1 scores for four zero-shot models stratified by BCA outcome group, evaluated on the pre-infusion CAR T product dataset. **b,** Fine-tuning effect on CD19-stimulated cells: dumbbell plot comparing zero-shot and fine-tuned weighted F1 for scGPT and scFoundation across BCA groups. **c,** Per cell-type zero-shot F1 scores averaged across BCA groups for all 17 CAR T cell subtypes. **d,** Heatmap of zero-shot F1 scores for three CD4+ subtypes averaged across BCA groups, highlighting consistent failure to annotate regulatory T cells across all models. **e,** Weighted F1 averaged across BCA groups under Basal and CD19-stimulated conditions for all six models. **f,** UMAP embeddings of pre-infusion product cells for ground truth and model-predicted cell type labels.

Interestingly, annotation performance was not uniform across clinical outcome groups. CAR T products from BCA-L patients with 8-year cancer-free remission consistently achieved the highest zero-shot accuracy (weighted F1 = 0.70–0.82), whereas the intermediate BCA2 group proved most difficult to annotate (weighted F1 = 0.58–0.68) (Fig. 2a). Although the underlying biological basis remains unclear, this observation suggests that infusion products associated with durable CAR T persistence retain transcriptional states that are more closely aligned with healthy immune-cell representations learned during pretraining, whereas products associated with intermediate persistence occupy a more heterogeneous or perturbed transcriptional landscape.

To determine whether this domain-shift penalty could be mitigated, we fine-tuned scGPT and scFoundation using the annotated CAR T reference dataset. Fine-tuning improved annotation accuracy for scGPT, while slightly decreasing accuracy for scFoundation (Fig. 2b). The gains exhibited by scGPT may reflect its relatively modest zero-shot performance and greater capacity for adaptation to the CAR T domain, whereas the lack of improvement exhibited by scFoundation are possibly due to its stronger pretrained baseline. Nevertheless, neither model fully recovered the performance observed on healthy PBMCs, indicating that domain adaptation alleviates, but does not eliminate, the challenges imposed by engineered cellular products.

We further evaluated the impact of antigen stimulation by comparing basal and CD19-stimulated CAR T infusion products. Across all four zero-shot models as well as two fine-tuned models, CD19 stimulation consistently reduced annotation accuracy by an additional 0.10–0.26 weighted F1 units relative to the basal condition (Fig. 2c). CD19 stimulation rapidly induces activation-associated transcriptional programs, including immediate-early response genes, cytokine signaling pathways and effector differentiation^26^, resulting in cellular states that are likely underrepresented in the predominantly resting immune cells used to pretrain current scFMs. These findings indicate that cellular activation introduces an additional layer of distribution shift beyond CAR T manufacturing itself.

Despite the overall reduction in annotation accuracy, performance varied markedly across individual CAR T cell populations. Several clinically important cell types consistently failed across all architectures. Most notably, CD4⁺ regulatory T (Treg) cells were essentially unidentifiable by every model (scGPT, F1 = 0.00; scFoundation, 0.03; CellPLM, 0.00; UCE, 0.00; Fig. 2d). Given the established role of Tregs in suppressing CAR T expansion and persistence, this systematic failure represents a potentially important limitation for clinical application. Other populations, including CD4⁺ immune-signaling, CD4⁺ basal FOS/JUN⁺ and MHC II⁺ cells, also exhibited consistently poor performance across models (Fig. 2e), suggesting that activated or transitional transcriptional states remain difficult for current scFMs to resolve. In contrast, abundant populations such as CD4⁺ memory, CD4⁺ activated FOS/JUN⁺, Th1/Tc1, proliferative G2M/S, CD8⁺ naïve and CD8⁺ XCL1/2⁺ cells were robustly identified by all architectures. Consistent with these quantitative results, UMAP visualization demonstrated that the global organization of the CAR T cellular landscape was largely preserved across models, while fine-tuning produced progressively sharper separation between neighboring cell populations (Fig. 2f).

### Model-derived cell compositions predict durable CAR T persistence despite imperfect cell annotation

The principal question of this study was not whether foundation models could perfectly annotate every individual cell, but whether they could recover sufficient biological information to predict clinically meaningful patient therapeutic outcomes. Previous studies have shown that the cellular composition of pre-infusion CAR T products influences *in vivo* expansion, persistence and therapeutic durability, with memory-like and less differentiated T-cell states generally associated with improved long-term responses. Because CAR T infusion products comprise heterogeneous mixtures of functionally distinct T-cell populations, we hypothesized that patient-level cell type composition inferred from scFM embeddings might preserve clinically informative signals even when individual cell labels are imperfect. We therefore calculated per-patient cell type proportions from the fine-tuned scGPT and scFoundation annotations and evaluated each population as an independent biomarker for the primary clinical endpoint: sustained long-duration BCA (BCA-L; n = 5) versus non-sustained BCA (combined BCA1 and BCA2; n = 28) under CD19 stimulation. Cell-type-specific discrimination was quantified using receiver operating characteristic (ROC) analysis, with statistical significance assessed by comparison against a 1,000-permutation label-shuffling null distribution (Fig. 3a). Here, we combined BCA1 and BCA2 into a single group, as AUC analysis showed that they were largely indistinguishable, with only one cell type showing significant discriminative power in the scFoundation model (Supplementary Fig. 1).

**Fig. 3:**
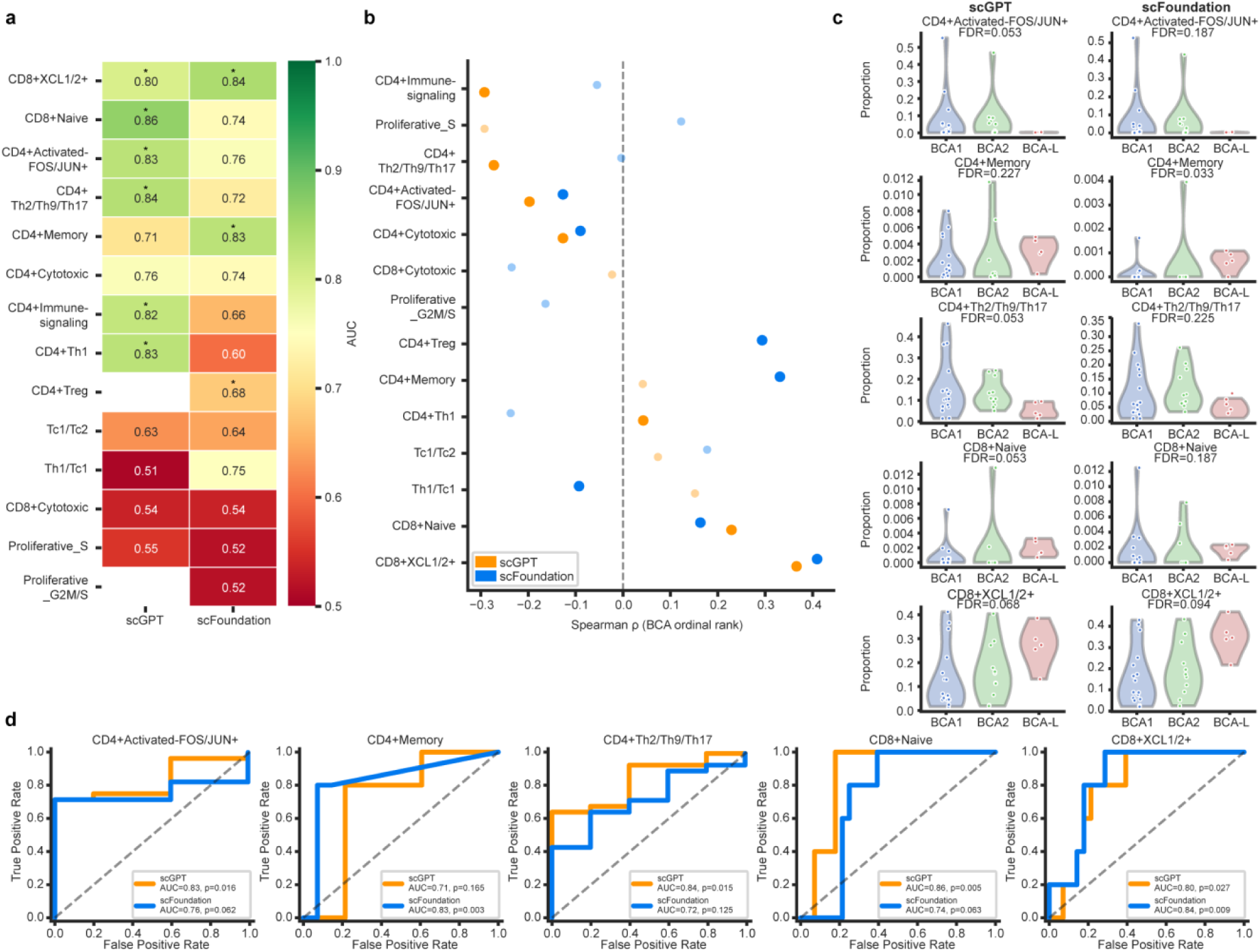
Clinical response prediction from model-derived cell type proportions in CD19-stimulated cells. a, AUC heatmap for discrimination of BCA-L from BCA1+BCA2 using cell type proportions derived from scGPT and scFoundation fine-tuned annotations under CD19 stimulation. Asterisks denote permutation-test significance (p<0.05). **b,** Spearman correlation between model-derived cell type proportions and ordinal BCA outcome rank under CD19 stimulation; filled points indicate Mann-Whitney FDR<0.2. **c,** Violin plots showing distributions of the five most discriminative cell type proportions across BCA groups for scGPT and scFoundation fine-tuned annotations under CD19 stimulation. **d,** ROC curves for BCA-L vs. BCA1+BCA2 classification using proportions of the five top-performing cell types under CD19 stimulation, for scGPT and scFoundation fine-tuned models.

Despite the reduction in per-cell annotation accuracy observed in the previous section, multiple scFM-derived cell populations retained strong predictive value for clinical outcome. Five cell types achieved permutation significance (P < 0.05) in at least one foundation model, including CD4⁺ activated-FOS/JUN⁺, CD4⁺ Th2/Th9/Th17, CD8⁺ naïve, CD8⁺ XCL1/2⁺ and CD4⁺ memory cells (Fig. 3a). Previous work^8^ reported that a Th2 functional phenotype is associated with durable responses in pediatric patients with B-ALL treated with CD19 CAR T cell therapy, broadly consistent with our observations. Among these cell types, CD8⁺ XCL1/2⁺ cells emerged as the only biomarker consistently identified by both scGPT and scFoundation (AUC = 0.80 and 0.84, respectively) under CD19 stimulation (Fig. 3a). This consistency distinguishes CD8⁺ XCL1/2⁺ cells from other candidate biomarkers whose significance depended on a particular model and identifies this population as the most reproducible predictor of durable CAR T persistence in our analysis. Given the established role of XCL1/XCL2 in recruiting XCR1⁺ cDC1 dendritic cells that support cytotoxic T-cell responses^27^, this association may reflect enhanced immune communication that promotes long-term CAR T persistence. More broadly, it suggests that durable clinical responses may depend not only on the intrinsic properties of CAR T cells but also on their ability to engage and coordinate with the host immune microenvironment.

To better understand the biological relationships underlying these associations, we next examined the correlation between inferred cell-type abundance and the ordinal spectrum of clinical response (BCA1, BCA2 and BCA-L). Cell populations associated with less differentiated and memory-like phenotypes, including CD8⁺ XCL1/2⁺, CD8⁺ naïve, Th1/Tc1 and CD4⁺ memory cells, showed progressively increasing abundance with longer CAR T persistence, whereas activated and proliferative populations, including CD4⁺ activated-FOS/JUN⁺, CD4⁺ Th2/Th9/Th17, proliferative S-phase and CD4⁺ immune-signaling cells, exhibited the opposite trend (Fig. 3b,c). These observations are broadly consistent with previous reports linking less differentiated T-cell states to prolonged CAR T persistence after infusion, while highly activated or terminally differentiated states are generally associated with reduced long-term functionality. We note, however, that the depletion of transcriptomically annotated CD4⁺ Th2/Th9/Th17 cells observed here should not be directly compared with the increased type-2 cytokine functionality previously reported for this cohort by Bai et al.^8^ as the former reflects embedding-based cell identity whereas the latter also measures functional cytokine output. We discuss this distinction further below.

Because antigen stimulation induces extensive transcriptional remodeling, we next asked whether biomarkers are more strongly depended on the stimulation state of the infusion product. Analysis of basal (unstimulated) samples identified a partially distinct set of predictive populations. Proliferative G2M/S and proliferative S-phase cells reached permutation significance in both scGPT and scFoundation, whereas CD8⁺ naïve cells remained predictive under both basal and stimulated conditions (Supplementary Fig. 2 and 3). Interestingly, greater abundance of proliferative populations before antigen stimulation was associated with shorter BCA, consistent with the possibility that highly proliferative products represent more differentiated cellular states with reduced capacity for sustained expansion following infusion and thus less durable response in patients.

Finally, we asked whether foundation-model-derived cell proportions provide information beyond expert cell annotations derived from the same scRNA-seq dataset. Repeating the biomarker analysis using the original manually curated cell labels demonstrated that scFM-derived proportions outperformed expert annotations for several cell populations (Supplementary Fig. 4). The largest improvements were observed for CD4⁺ Th2/Th9/Th17 cells (AUC = 0.84 versus 0.59; ΔAUC = +0.25) and CD4⁺ memory cells (AUC = 0.83 versus 0.69; ΔAUC = +0.14), whereas performance for CD8⁺ XCL1/2⁺ cells was comparable between the two approaches. Because both analyses originate from the same underlying sequencing data, this comparison does not constitute independent biological validation. Instead, it suggests that foundation-model embeddings capture continuous transcriptional variation that may be partially obscured when cells are assigned to discrete expert-defined categories. Consistent with this interpretation, no cell population achieved statistical significance in the negative-control comparison between BCA1 and BCA2 (all P ≥ 0.05), indicating that the predictive signal recovered by scFMs is specific to the biologically distinct phenotype of sustained long-term CAR T persistence rather than reflecting nonspecific inter-patient variability.

### Cross-context generalization to post-infusion CAR T cells is architecture-dependent

A key question for scFMs to predict clinical response is whether representations learned from healthy immune cells and stimulation or activation of CAR T infusion products generalize to CAR T cells functioning in patients after infusion. Following infusion, CAR T cells undergo extensive transcriptional remodeling in response to antigen encounter, proliferation, immune activation, and progressive differentiation, generating cellular states that differ substantially from both healthy PBMCs and the manufactured infusion product^28^. To examine this challenging setting, we evaluated all four zero-shot foundation models on a published post-infusion peripheral blood scRNA-seq dataset comprising 9,525 CAR T cells collected from a patient following BCMA CAR T therapy. Because this dataset represents a single patient, we consider the analysis a proof-of-concept assessment of cross-context generalization rather than a cohort-level evaluation.

Across all four models, we observed a common failure mode involving clinically important T-cell states. CD8⁺ exhausted T cells, CD8⁺ CD160⁺ T cells, CD4⁺CD25⁺FOXP3⁺ Tregs and CD4⁺CD25⁺ Tregs were uniformly annotated with near-zero F1 scores regardless of architecture (Fig. 4a). Among these populations, exhausted CD8⁺ T cells are of particular biological importance because progressive T-cell exhaustion is a hallmark of CAR T dysfunction and represents one of the major determinants of treatment failure. Their consistent absence from foundation-model predictions therefore suggests that dysfunctional and chronically stimulated T-cell states remain poorly represented in current pretraining datasets, which are composed predominantly of healthy immune cells. In contrast, the dominant population in this dataset, CD8⁺ JUN⁺CD69⁺ memory-like T cells, accounting for approximately 73% of all cells, was accurately identified across all models, illustrating that aggregate performance metrics can remain relatively high even while clinically critical rare populations are systematically missed.

**Fig. 4:**
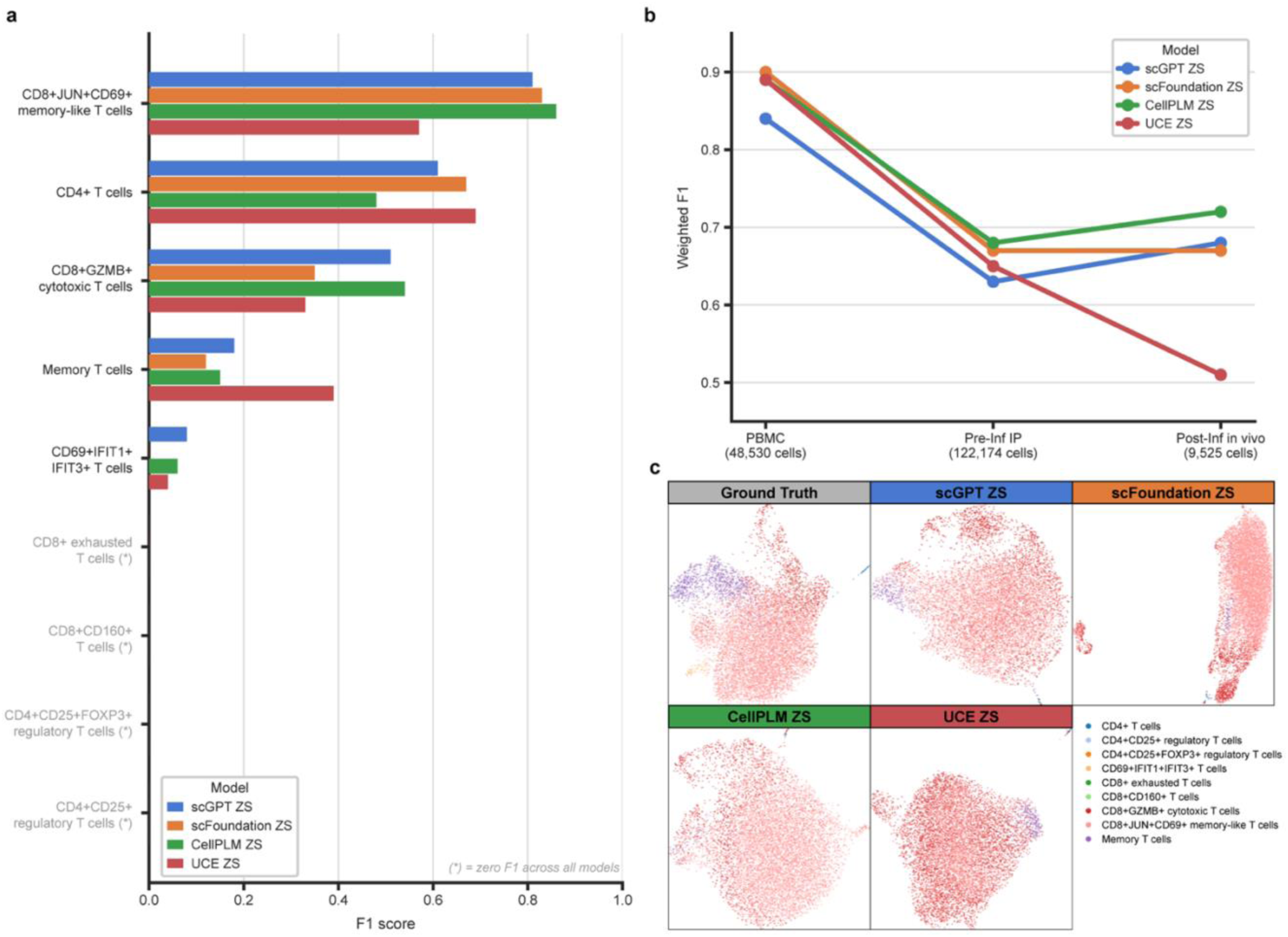
Cross-context generalization and post-infusion *in vivo* annotation performance. a, Per cell-type zero-shot F1 scores for nine T cell subtypes in the post-infusion *in vivo* dataset. Cell types with zero F1 across all models are shown in gray. **b,** Weighted F1 scores across three evaluation contexts (PBMC, BCA1 cohort of pre-infusion CAR T product, post-infusion *in vivo*) for four zero-shot models. **c,** UMAP embeddings of post-infusion cells colored by ground truth and model-predicted cell type labels for scGPT zero-shot, scFoundation zero-shot, CellPLM zero-shot, and UCE zero-shot.

Although these shared failure modes were consistent across models, overall generalization performance differed substantially between architectures (Fig. 4b). CellPLM achieved the highest weighted F1 score on the post-infusion dataset (0.72), despite not consistently outperforming other models on the PBMC or pre-infusion CAR T benchmarks. scGPT and scFoundation exhibited comparable performance (both approximately 0.68), whereas UCE showed a pronounced decline from its strong performance on healthy PBMCs (weighted F1 = 0.89) to only 0.51 on post-infusion CAR T cells. Consistent with this quantitative result, UMAP visualization revealed that UCE assigned the majority of cells to a single dominant population, indicating a loss of resolution under this challenging distribution shift (Fig. 4c). Collectively, these findings demonstrate that although all current scFMs experience reduced performance when applied to post-infusion CAR T biology, the extent of degradation is strongly architecture-dependent. More broadly, they underscore that robust performance on healthy reference datasets does not necessarily predict reliable behavior in clinically relevant disease settings, highlighting the need for context-specific fine turning and cross-context generalization as a critical benchmark for future generations of scFMs.

### Clinical decision-support AI agent predicts durable CAR T treatment response

Having established that scFM-derived cell compositions capture clinically relevant information despite imperfect cell-level annotation, we next asked whether these predictions could be translated into an interpretable clinical decision-support tool for individualized outcome prediction. We therefore developed a locally deployable clinical decision-support AI agent that integrates foundation model-based cell-state profiling with patient-level risk prediction to estimate the probability of sustained long-duration BCA from a patient’s pre-infusion CAR T product (Fig. 5a). The agent automates a workflow that would otherwise require bioinformatics experts to execute a series of sophisticated and often labor-intense computational analysis steps, including cell-state annotation, quantification of cellular composition, biomarker interpretation and statistical clinical outcome prediction, thereby providing an accessible framework for translating scFMs into clinically actionable predictions. We selected the fine-tuned scGPT model as the backbone of the current implementation because it achieved the highest annotation accuracy under CD19 stimulation (Fig. 2b), although systematic comparison between zero-shot and fine-tuned models for clinical prediction remains an important direction for future investigation.

**Fig. 5:**
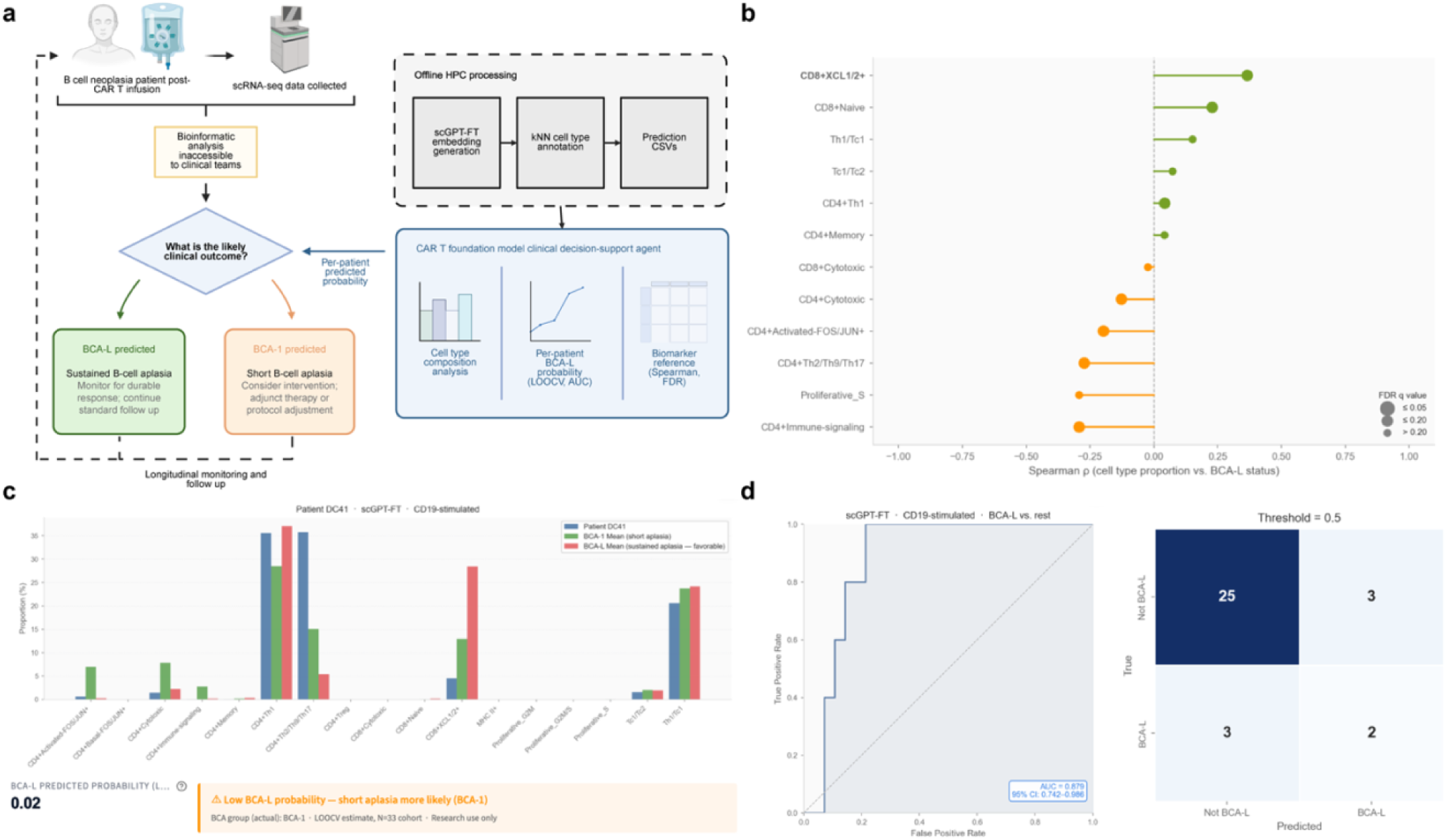
CAR T foundation model clinical decision-support agent for B-cell aplasia outcome prediction. a, Clinical decision framework and system architecture of the CAR T FM clinical decision-support agent. scGPT-FT cell type predictions are generated offline via HPC and loaded into a locally deployable interface that returns a per-patient BCA-L predicted probability to inform clinical decision-making. **b,** Spearman correlation (ρ) between scGPT-FT predicted cell type proportions and BCA-L status (n=33, CD19-stimulated). Dot size reflects FDR-adjusted q value (Benjamini-Hochberg; 17 cell types). Green: enriched in BCA-L; orange: depleted in BCA-L. **c,** Illustrative representative interface output for an example BCA-1 patient, showing predicted cell type composition relative to BCA-1 and BCA-L cohort means and a BCA-L predicted probability of 0.02. **d,** LOOCV classification performance of the BCA-L vs. rest logistic regression classifier (scGPT-FT, CD19-stimulated; n=33). AUC = 0.879 (95% CI: 0.742–0.986, bootstrap n=1,000 iterations). Confusion matrix shows predicted vs. true BCA-L vs. non-BCA-L classification at threshold = 0.5. Created in BioRender. Shen, L. (2026) https://BioRender.com/vu0p0nq is licensed under CC BY 4.0.

The decision-support workflow consists of two sequential stages. First, single-cell transcriptomes from the pre-infusion CAR T product are processed by the foundation model to generate cell embeddings and infer cell identities, from which patient-specific cell type composition is calculated. These compositional features are subsequently used by a logistic regression classifier to estimate the probability of durable CAR T persistence, while simultaneously providing an interpretable summary of the cellular populations contributing to the prediction. In the current prototype, foundation model inference is performed offline and the resulting patient-level composition profiles are loaded into the application to ensure complete reproducibility of the reported analyses. In a future clinical implementation, this inference step could be performed directly on newly generated patient scRNA-seq data, enabling fully automated prediction at the point of care.

The biological features contributing to clinical prediction closely mirrored the biomarker analyses described above (Fig. 5b). Cell populations enriched in patients with durable CAR T persistence, including CD8⁺ XCL1/2⁺, CD8⁺ naïve, Th1/Tc1, CD4⁺ Th1 and CD4⁺ memory cells, contributed positively to the predicted probability of sustained BCA. Conversely, activated and proliferative populations, including CD4⁺ activated-FOS/JUN⁺, CD4⁺ Th2/Th9/Th17, proliferative S-phase and CD4⁺ immune-signaling cells, were associated with reduced predicted persistence. Figure 5c illustrates a representative analysis of a BCA1 patient, for whom the agent predicted a probability of only 0.02 for long-duration BCA, consistent with the observed clinical outcome.

To evaluate the predictive performance, we performed leave-one-out cross-validation across all 33 patients using CD19-stimulated infusion products. The resulting classifier achieved an AUC of 0.879 (95% bootstrap confidence interval, 0.742–0.986; 1,000 bootstrap resamples), demonstrating that scFM-derived cell composition analysis using an AI agent enables accurate patient-level prediction despite imperfect annotation at the individual-cell level (Fig. 5d). At a decision threshold of 0.5, the classifier correctly identified 25 of 28 patients without sustained BCA and 2 of 5 patients with long-duration persistence. Using basal (unstimulated) samples yielded a slightly lower AUC of 0.850, supporting CD19 stimulation as the preferred input for clinical prediction. Although these results are encouraging, the limited number of patients with sustained long-duration BCA (n = 5) results in relatively wide confidence intervals. Consequently, the present decision-support agent should be viewed as a proof-of-concept demonstrating the feasibility of foundation model-guided clinical prediction, with prospective validation in larger multi-center cohorts required before routine clinical deployment.

## Discussion

The potential of CAR T cell therapy in B-ALL is clearly limited by response durability, but the pre-infusion product offers a uniquely accessible window into the determinants of long-term persistence. Our central finding is that scFM embeddings carry sufficient phenotypic signal in the pre-infusion product to stratify long-duration from shorter-duration outcomes at leave-one-out cross-validated AUC of 0.879, despite a 0.21–0.25 weighted F1 drop on per-cell annotation relative to a healthy PBMC reference. Cohort-level cell type proportions seem to average out much of the residual annotation error, preserving the downstream clinical signal. CD8^+^XCL1/2^+^ cells emerged as the only biomarker reaching permutation significance across both scGPT and scFoundation under CD19 stimulation, identifying a candidate marker of sustained CAR T persistence. XCL1 and XCL2 chemokines recruit XCR1^+^ cross-presenting dendritic cells and amplify adaptive immune responses^29, 30^, providing a plausible mechanistic explanation for the observed enrichment in long-duration BCA patients. That foundation-model-derived proportions exceeded ground-truth-derived proportions on multiple discriminative features, with ΔAUC up to +0.25 for CD4^+^Th2/Th9/Th17, suggesting scFM embeddings capture continuous variation in cell state that is partially obscured by discrete categorical labeling. However, this comparison is a within-dataset benchmark and orthogonal validation remains essential.

Our cell-label-level findings on the CD4^+^Th2/Th9/Th17 compartment introduce nuance to the prior characterization of this same cohort by Bai et al.^8^, which reported that elevated type-2 functionality in the pre-infusion product is associated with eight-year sustained leukemia remission. This cohort was previously characterized by the same group^31^ in a smaller subset analysis, where deficiency in Th2 functional output in CAR T cells was associated with CD19-positive relapse. We observe that cells assigned the CD4^+^Th2/Th9/Th17 transcriptomic label by foundation-model-derived annotation are depleted in BCA-L, which on its face appears to invert the published direction of association. We do not interpret these as contradictory findings. Bai et al. measured type-2 functionality through cytokine functional output, quantified by intracellular cytokine staining, multiplexed secretomic assay, and a transcriptomic type-2 score derived from gene-set scoring, with orthogonal validation by IL-4 supplementation experiments. Our measurement is the proportion of cells whose embedding-space neighborhood maps to a CD4^+^Th2/Th9/Th17 reference cluster under k-NN reference mapping. These are related but distinct—a cell may carry elevated type-2 cytokine output without occupying the embedding neighborhood of canonically labeled Th2/Th9/Th17 reference cells, particularly in an engineered and antigen-stimulated CAR T product where cytokine programs and lineage-defining transcriptomic patterns may decouple. A complete reconciliation would require co-registering scFM-derived per-cell embeddings with paired cytokine measurements at the single-cell level, which our data does not currently support. The cell-label-level Th2/Th9/Th17 finding reported in this study is hypothesis-generating and complementary to, rather than displacing our previous functional characterization.

A growing benchmarking literature has reported that current scFMs do not consistently outperform classical methods on standard zero-shot tasks, particularly under distribution shift^20–22^. Kedzierska et al.^20^ find that scFMs frequently underperform simple PCA baselines on zero-shot integration and label transfer across multiple atlases. Boiarsky et al.^21^ demonstrate that scGPT’s zero-shot embeddings can be matched or exceeded by carefully tuned classical pipelines on standard cell-typing benchmarks. Liu et al.^22^ report broadly similar findings across multiple architectures and tasks. Our results are largely consistent with this critique on its own terms: PBMC zero-shot performance approaches a performance ceiling that classical reference-mapping methods also reach^32, 33^, and the engineered CAR T product imposes a domain-shift penalty that fine-tuning only partially recovers. However, while the published critique literatures evaluate scFMs on per-cell annotation accuracy in standard benchmark settings, we instead evaluate cohort-aggregated cell type proportions on a downstream clinical inference task. These are different evaluation regimes, and they yield different conclusions. On per-cell annotation in standard settings, scFMs are not transformatively better than classical methods. On cohort-aggregated phenotypic signal in a clinically meaningful task, scFM-derived features are sufficient to support stratification at LOOCV AUC of 0.879. We do not view these findings as in tension. Together, the literature and our results jointly suggest that the right test for clinical translatability of scFMs is whether their representations carry phenotype-relevant signal in real downstream scenarios such as domain shift, instead of their performance on standard integration benchmarks (e.g., scVI-style batch-integration tasks). Three qualifications matter in this situation: the strongest clinical performance is achieved only after fine-tuning, the signal is recoverable at the level of cohort-aggregated proportions rather than per-cell labels, and the signal is concentrated in the extreme BCA-L contrast (the BCA1-versus-BCA2 negative control yielded no permutation-significant features for either model). The deployable use case implied is biomarker discovery and cohort-scale risk stratification rather than deterministic per-patient classification.

Two failures surfaced across all models, and both negatively impact potential for clinical use. CD4^+^ Treg cells were annotated at near-zero F1 by every model in every context we evaluated, including the PBMC reference, the pre-infusion product, and the post-infusion peripheral blood. Tregs are central modulators of CAR T efficacy and persistence, and their consistent absence from scFM-derived annotations represents a systematic gap that fine-tuning on T-cell-rich reference data does not correct. Similarly, post-infusion exhausted CD8^+^ subsets, which are the canonical markers of CAR T dysfunction, failed across all four models on the post-infusion dataset. Both failures likely reflect underrepresentation of regulatory and disease-activated immune states in the pretraining corpora that anchor current scFMs. This is primarily a pretraining corpus design problem, and it argues for deliberate enrichment of disease-activated, exhausted, and regulatory immune states in next-generation scFM training data before clinical deployment in CAR T or related cellular therapy contexts.

A second concern with direct deployment implications is the partial cross-architecture agreement on biomarker identity. While CD8^+^XCL1/2^+^ achieved cross-model significance, several other discriminative cell types reached permutation significance for one model but not the other (CD4^+^memory significant for scFoundation but not scGPT; CD8^+^naive significant for scGPT but not scFoundation), and the cross-context generalization behavior of UCE, which was strong on PBMC but collapsed on the post-infusion dataset to weighted F1 = 0.51, illustrates how dramatically architecture choice can shape downstream results. We note that this evaluation used the 4-layer UCE variant; whether the 33-layer flagship model would exhibit the same post-infusion failure is an open question that would substantially affect interpretation of UCE-specific findings. For a clinical decision-support agent deployed prospectively, this means the choice of foundation model is a clinically relevant decision instead of solely methodological, and argues for ensemble approaches, orthogonal validation against non-transcriptomic biomarkers, or both, before any prospective deployment.

This study has several limitations that scope the strength of our claims. The BCA-L cohort is small (n=5), reflecting the inherent rarity of sustained eight-year relapse-free BCA in prospectively collected pediatric CAR T datasets. The resulting wide bootstrap confidence interval (AUC 95% CI 0.742–0.986) also underscores the need for prospective multicenter validation. Our post-infusion generalization analysis was limited to a single publicly available patient profiled in the context of BCMA-targeted CAR T therapy for plasma cell leukemia^24^; this enabled a proof-of-concept cross-context evaluation, but the findings should not be interpreted as direct evidence of generalization to CD19-directed CAR T or to larger post-infusion cohorts. The expert ground-truth annotations against which scFM-derived proportions were compared were derived from the same scRNA-seq data, and orthogonal validation by flow cytometry or multiplexed secretomic profiling would be required to distinguish whether the foundation-model-exceeds-ground-truth signal reflects genuine capture of continuous cell state variation or correlated noise structure between label assignments. We selected scGPT-FT as the agent backbone on annotation performance under CD19 stimulation; whether fine-tuning additionally improves clinical discrimination over zero-shot inference remains open. Finally, the agent does not currently quantify per-cell model uncertainty, and cells near scFM decision boundaries are silently misannotated.

Our findings carry a broader implication for the evaluation of scFMs for clinical translation. The current benchmarking convention privileges per-cell annotation accuracy on standardized reference datasets, and conclusions about scFM utility are typically drawn from these aggregate metrics. Our work suggests two amendments. First, evaluation should be downstream-task-aware: a model that is unimpressive on per-cell annotation may still carry decisive signal for cohort-level clinical inference, and the converse may also hold. Second, evaluation should be cross-architecture by design rather than as a default convenience. The partial agreement on biomarker identity we observe between scGPT and scFoundation, and the catastrophic divergence we observe for UCE on the post-infusion dataset, are exactly the kinds of findings that single-architecture evaluations cannot surface but that any clinically deployed scFM will encounter. As scFMs move toward deployment in cellular therapy and other translational settings, evaluation frameworks that combine task-relevant downstream metrics with cross-architecture comparison provide a more thorough picture of clinical readiness than per-cell accuracy alone.

Several extensions would substantively extend this work. The most important is prospective validation of CD8^+^XCL1/2^+^ as a biomarker of sustained CAR T persistence in an independent multicenter cohort, ideally with paired flow cytometric or secretomic confirmation. A fully integrated point-of-care agent that performs scFM inference on each new patient’s pre-infusion product on GPU-backed compute, rather than relying on offline pre-computed CSV files, is a near-term engineering target. Comparison of fine-tuned versus zero-shot scFM inputs to the clinical classifier would clarify whether fine-tuning improves clinical signal or only annotation accuracy. Ensemble approaches that aggregate proportional features across multiple foundation model architectures may both improve cross-cohort robustness and surface architecture-specific biomarker signals that a single-model agent would miss. More broadly, the systematic Treg and exhausted CD8^+^ failures we observe argue for the deliberate inclusion of regulatory and disease-activated immune states in next-generation pretraining corpora, particularly for any scFM intended for cellular therapy applications.

## Methods

### Datasets and BCA outcome group definitions

Three publicly available single-cell RNA sequencing datasets anchored the analysis. The pre-infusion CAR T cell product cohort comprised 231,088 cells from 33 pediatric patients with B-cell acute lymphoblastic leukemia enrolled across the first two CD19-CAR T trials at the Children’s Hospital of Philadelphia, originally published by Bai et al.^8^ (NCBI GEO accession GSE262072), with each product profiled under both basal (unstimulated) and CD19-stimulated conditions. Patients were stratified into three mutually exclusive outcome groups by duration of post-infusion B-cell aplasia (BCA): BCA1 (short-duration aplasia, n=17), BCA2 (intermediate-duration, n=11), and BCA-L (long-duration sustained aplasia, n=5). The cohort labeled BCA3 in the original Bai 2024 nomenclature corresponds to the BCA2 cohort used here; this label normalization was applied uniformly during data ingestion. A reference partition of healthy normal donor cells, distinct from the patient cohort used for clinical prediction, was used for fine-tuning and k-NN annotation. The healthy peripheral blood mononuclear cell (PBMC) reference comprised 48,530 cells from 8 donors with 31 annotated cell types (Hao et al.^23^, GEO accession GSE164378), and may be present in the pretraining data of the evaluated models. The post-infusion peripheral blood dataset comprised 9,525 cells from a single patient profiled at the remission timepoint following BCMA-targeted CAR T therapy for plasma cell leukemia (Li et al.^24^); the corresponding peak-phase cells from the same patient served as the reference partition for k-NN annotation. All input data were AnnData HDF5 files (.h5ad).

### Computational environment

All computations were performed on the Yale McCleary high-performance computing cluster. GPU-accelerated jobs were submitted via SLURM, with UCE embedding generation allocated 1 GPU, 8 CPUs, 64 GB RAM, and a 7-hour wall-time per job. Four conda environments were maintained to manage dependency conflicts between models: scGPT (Python 3.9, PyTorch 2.1.2, CUDA 11.8, flash-attn<1.0.5), scFoundation (Python 3.9, PyTorch 2.1.2, CUDA 11.8, local-attention, einops), CellPLM (PyTorch 1.13.0+cu117, faiss-gpu, CellPLM), and UCE (Python 3.10). FAISS-based reference mapping for UCE embeddings was performed in the scFoundation environment due to FAISS dependency incompatibility in the UCE environment.

### Foundation models

Four single-cell RNA sequencing foundation models were evaluated. All four were assessed in zero-shot configuration; scGPT and scFoundation were additionally fine-tuned on the CAR T reference partition. *scGPT* (Cui et al.^13^) is a generative pretrained transformer with embedding dimension 512, 12 transformer layers, 12 attention heads, and a 60,697-gene vocabulary, pretrained on 33 million normal human cells from the CELLxGENE Census; the pretrained whole-human checkpoint (best_model.pt; ∼53 million parameters^34^) was used. *scFoundation* (Hao et al.^14^) implements a masked autoencoder with automatic expression binning (MaeAutobin); the 100-million-parameter encoder uses hidden dimension 768, 12 layers, and 12 attention heads, with a 19,264-gene index, pretrained on over 50 million human single-cell transcriptomes. *CellPLM* (Wen et al.^15^) is a cell-as-token spatial transformer pretrained on >10 million cells spanning scRNA-seq and spatially resolved transcriptomic profiles; the 85-million-parameter checkpoint (20231027_85M) was used in zero-shot mode (the public API does not expose fine-tuning). *UCE* (Rosen et al.^16^) generates universal cell embeddings using ESM2 protein language model embeddings as gene tokens, pretrained on approximately 36 million cells across eight species (the majority derived from CELLxGENE Census). The published flagship UCE model is a 33-layer transformer with approximately 650 million parameters; we evaluated the smaller 4-layer model checkpoint (4layer_model.torch), which is distributed alongside the flagship model as the default option in the official UCE inference pipeline and is the variant most evaluated in published scFM benchmarking studies. Human was specified as the target species.

### Data preprocessing

Each model received raw count matrices passed through model-specific preprocessing to align gene vocabularies and tokenize expression. For scGPT, the scgpt.preprocess.Preprocessor class was applied with total-count normalization to 10⁴, 51 expression bins, no filtering, and tokenization with append_cls=True and max_seq_len=3001. For scFoundation, dataset columns were reordered to match the 19,264-gene index, padding missing genes with zeros; for fine-tuning, the top 5,000 highly variable genes (Seurat v3 flavor) intersected with the scFoundation index were retained. CellPLM required Ensembl ID gene identifiers, with symbol-to-Ensembl conversion performed via the biothings API. UCE accepted raw counts directly with --allow_non_counts, performing normalization internally. Detailed gene-recovery statistics across models are provided in Supplementary Methods.

### Fine-tuning

Both scGPT and scFoundation were fine-tuned on the healthy normal donor partition dataset using a patient-level train/val split. scGPT was fine-tuned for 10 epochs with batch size 64, learning rate 1×10⁻⁴, dropout 0.2, AMP enabled, flash attention enabled, and the CLS classification objective active; pretrained architecture parameters (embsize=512, nlayers=12, nheads=12) were preserved during fine-tuning. scFoundation was fine-tuned via a custom FineTuneMaeCell wrapper appending a BatchNorm → Linear → ReLU → Linear classification head to the encoder output, trained for 5 epochs with batch size 64, gradient accumulation 16, AdamW optimizer with differential learning rates (1×10⁻⁴ for backbone, 1×10⁻³ for classification head, weight decay 1×10⁻² for both), and CrossEntropyLoss. Cell embeddings for both models were extracted from the CLS token (scGPT) or via max-pooling over the encoder sequence dimension (scFoundation). scFoundation fine-tuning used a custom implementation as no official fine-tuning script is provided by the model authors; this is documented as a source of experimental variance relative to scGPT fine-tuning. Fine-tuned checkpoints achieved validation accuracy of 0.872 (scGPT) and 0.878 (scFoundation, epoch 5). CellPLM and UCE were evaluated zero-shot only.

### FAISS-based reference mapping

A unified k-nearest-neighbor reference mapping protocol was applied across all four models for cell type annotation. Reference cell embeddings were indexed using a GPU-accelerated FAISS IndexFlatL2 structure performing exact nearest-neighbor search under squared Euclidean distance. Query cells were assigned cell type labels by majority vote across their k=11 nearest reference neighbors; ties were broken by selecting the label with the smallest mean L2 distance to the query cell. For pre-infusion product annotation, the reference is the healthy normal donor partition dataset; for post-infusion *in vivo* annotation, the reference was the peak-phase partition of the Li et al.^24^ dataset. Classification performance was evaluated using sklearn.metrics.classification_report with weighted F1 as the primary aggregate metric.

### Clinical prediction analysis

For each model and stimulation condition, prediction CSVs from the three BCA groups were concatenated into a unified per-patient cell type proportion matrix (33 patients × cell types, normalized to row sum = 1.0). Univariate biomarker analysis was performed for the primary clinical contrast (BCA-L vs. BCA1+BCA2, n=5 vs. n=28) under CD19 stimulation, with the basal condition evaluated as a secondary analysis. Receiver operating characteristic area under the curve (AUC) was computed per cell type using sklearn.metrics.roc_auc_score, with values reflected to the [0.5, 1.0] range to capture discrimination magnitude regardless of effect direction. Statistical significance of each AUC was assessed by two-sided permutation test with 1,000 BCA-L label-shuffle iterations (RANDOM_SEED=42), with p-value defined as the proportion of permutations yielding |AUC_perm − 0.5| ≥ |AUC_obs − 0.5|. Univariate AUC values were computed on the full 33-patient cohort as a screening metric for biomarker identification, not as cross-validated estimates of generalization performance. Spearman rank correlation between cell type proportions and ordinal BCA rank (BCA1=1, BCA2=2, BCA-L=3) was computed across all 33 patients; Mann-Whitney U tests (two-sided) compared proportions between BCA outcome groups. Multiple testing correction across the cell types tested within each comparison used the Benjamini-Hochberg false discovery rate procedure. The BCA1-vs-BCA2 contrast (n=17 vs. n=11) was evaluated as a negative control; only CD4+Th1 reached nominal permutation significance for scFoundation in this comparison, consistent with the false-positive rate expected across this number of tests at p<0.05.

### Clinical decision-support agent

The clinical decision-support agent was implemented as a locally deployable Streamlit web application with three pages (Patient Analysis, Cohort LOOCV, Biomarker Reference) consuming six pre-computed scGPT-FT prediction CSV files (3 BCA groups × 2 stimulation conditions). Foundation model inference (embedding generation and k-NN annotation) was performed offline on HPC infrastructure, with per-cell predictions written to disk as CSV files; the agent loads these files at runtime and performs only downstream classification and visualization. This pre-computed-input architecture was adopted to fix the inference substrate across all reported analyses and to permit auditability of the prediction pipeline without re-running embedding generation. Consequences of this architectural choice include no GPU requirement at runtime, no internet dependency, and complete reproducibility of the reported figures from the deposited CSV files alone; these properties support manuscript-level reproducibility rather than constituting features of an intended clinical deployment, which would require GPU-backed scFM inference on each new patient’s product. A leave-one-out cross-validation logistic regression classifier (L2 regularization, C=1.0, lbfgs solver, max_iter=1000) was trained on per-patient scGPT-FT-derived cell type proportions under CD19 stimulation, with features standardized by StandardScaler fit on the training fold at each LOOCV iteration. BCA2 patients were grouped with BCA1 in the non-BCA-L reference class, reflecting the clinically actionable distinction between sustained and non-sustained aplasia; a three-class formulation was underpowered at n=11 for BCA2. Bootstrap 95% confidence intervals for the LOOCV AUC were computed from 1,000 bootstrap resamples of the LOOCV predicted probabilities and true labels (random seed = 42), with the interval defined by the 2.5th and 97.5th percentiles of the bootstrap AUC distribution^35^.

### Statistical analysis and visualization

All statistical analyses were performed in Python 3.9 using scipy.stats (Mann-Whitney U, Spearman ρ), sklearn.metrics (ROC-AUC, classification_report), and statsmodels.stats.multitest (Benjamini-Hochberg FDR^36^). Annotation accuracy was reported using weighted F1 as the primary aggregate metric, accounting for class imbalance; per-class F1 scores were retained for assessment of rare-cell-type performance. Given the rare long-duration group (n=5 BCA-L), AUC-based biomarker associations were assessed by permutation test and reported as nominal p-values, and are interpreted as exploratory rather than confirmatory, consistent with the primary study of this cohort. For the proportion-level and rank-correlation analyses, Benjamini–Hochberg q-values were computed across the cell types tested within each comparison and are shown in the corresponding figures. Figures were generated with matplotlib and seaborn using Arial typography. The random seed for permutation testing and stochastic analyses was set to 42 throughout.

## Data availability

This study did not generate new experimental data. The healthy human PBMC dataset by CITE-seq was from GSE164378. The pre-infusion CAR T infusion product dataset by combined scRNA-seq and CITE-seq was from GSE262072. The human post-infusion *in vivo* dataset by scRNA-seq was from GSE151310. Source data for figures and analysis are provided with this paper.

## Code availability

All foundation model checkpoints were obtained from each model’s published GitHub repository: [scGPT](https://github.com/bowang-lab/scGPT), [scFoundation](https://github.com/biomap-research/scFoundation), [CellPLM](https://github.com/OmicsML/CellPLM), and [UCE](https://github.com/snap-stanford/UCE). The source code for the clinical decision-support agent, together with the reference prediction tables it utilizes, is available in the GitHub repository at https://github.com/klshen8386/carT-clinical-agent and archived at https://doi.org/10.5281/zenodo.20389684. The model evaluation pipeline (preprocessing, fine-tuning, embedding generation, and reference mapping) together with the resulting per-cell prediction tables and per-patient cell type proportion tables, is available at https://doi.org/10.5281/zenodo.20389574.

## Acknowledgements

Computational data analysis was conducted with Yale High Performance Computing clusters (HPC). We acknowledge the support received from the SU2C Convergence 2.0 Grant. Cartoons in Figures 1 and 5 were created with BioRender.com. This article reflects the views of the authors and should not be construed as representing the views or policies of the institutions that provided funding.

## Author contributions

L.S. conceived the foundation model evaluation framework, performed the cross-architecture benchmarking, designed and implemented the clinical decision-support agent, conducted all statistical analyses, generated the figures, and wrote the manuscript. Z.B. provided the patient cohort scRNA-seq data and B-cell aplasia outcome annotations from the published study cohort^8^, contributed expertise in CAR T biology and infusion-product characterization, and provided manuscript revision. M.Y. provided computational infrastructure support, contributed to pipeline implementation and debugging, and provided manuscript revision. N.L. provided guidance on AI agent design and provided manuscript revision. R.F. supervised the project, contributed to conceptual framing and clinical translational direction, data interpretation, and manuscript revision. All authors reviewed and approved the final manuscript.

## Supplementary Information

**Supplementary Fig. 1:**
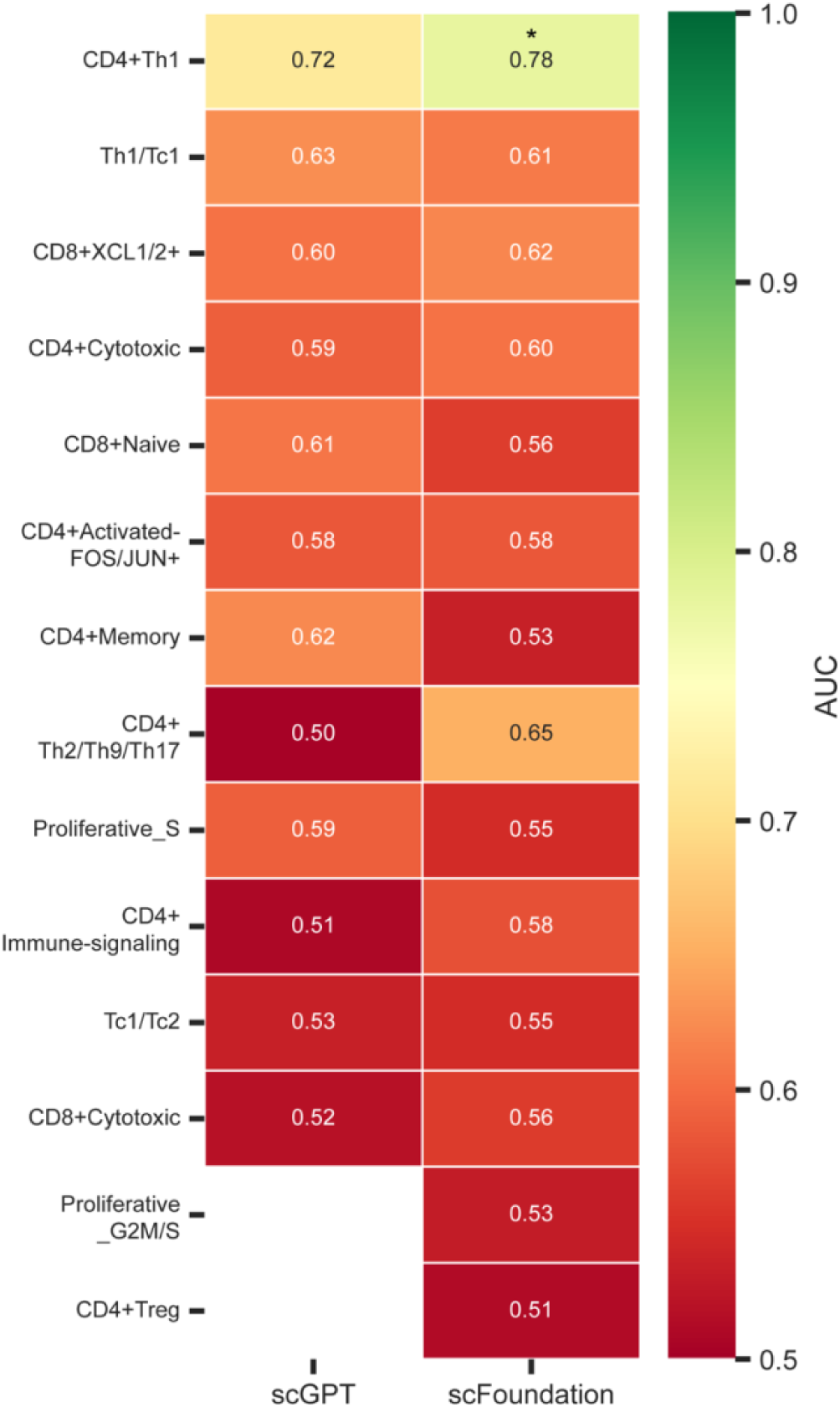
AUC heatmap for discrimination between BCA1 and BCA2 using cell type proportions derived from scGPT and scFoundation fine-tuned annotations under CD19 stimulation condition. Asterisks denote permutation-test significance (p<0.05).

**Supplementary Fig. 2:**
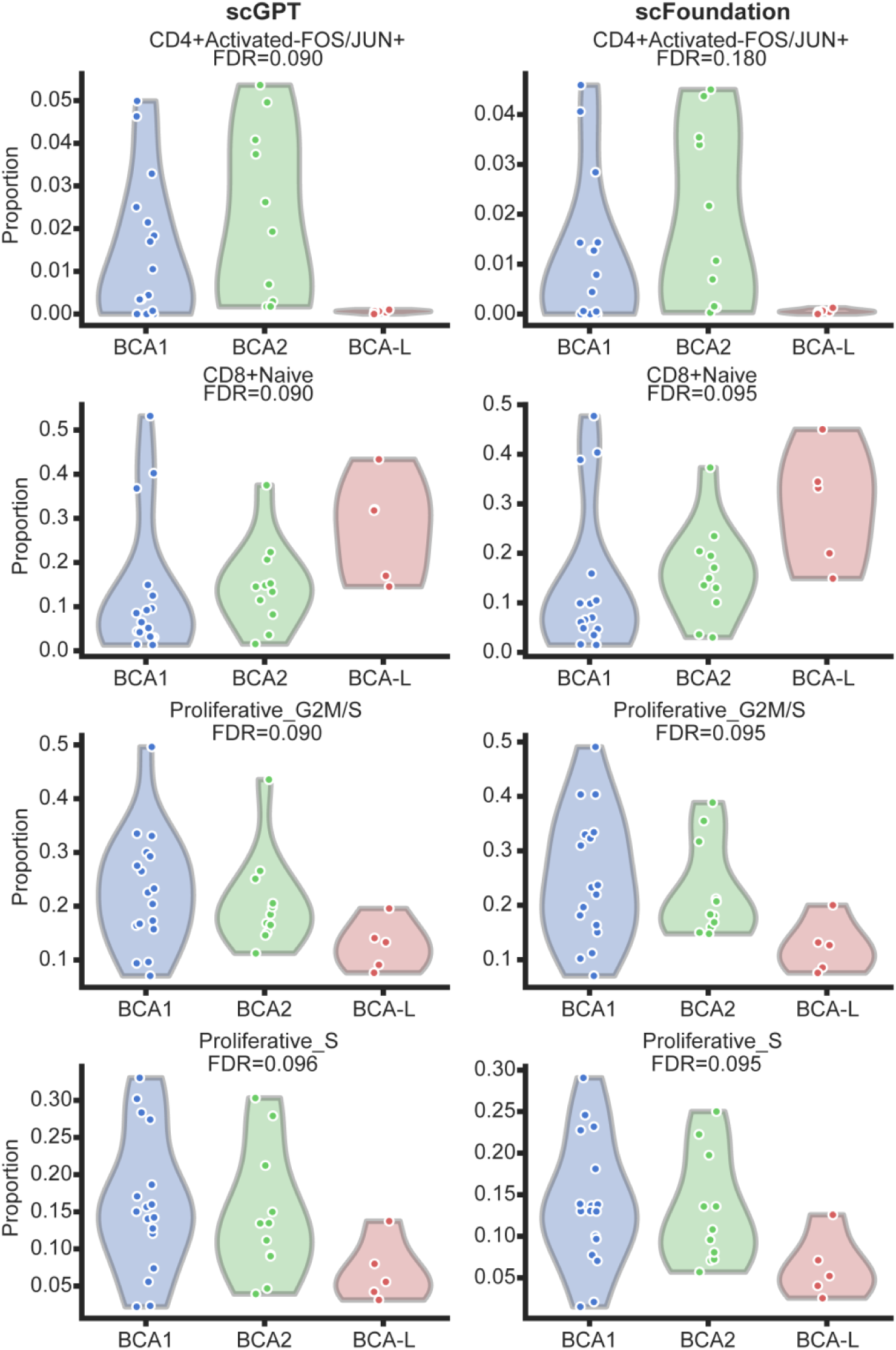
Violin plots showing distributions of the four most discriminative cell type proportions across BCA groups (BCA1, BCA2, BCA-L) for scGPT and scFoundation fine-tuned annotations under basal (unstimulated) condition; FDR values from Mann-Whitney tests are shown.

**Supplementary Fig. 3:**
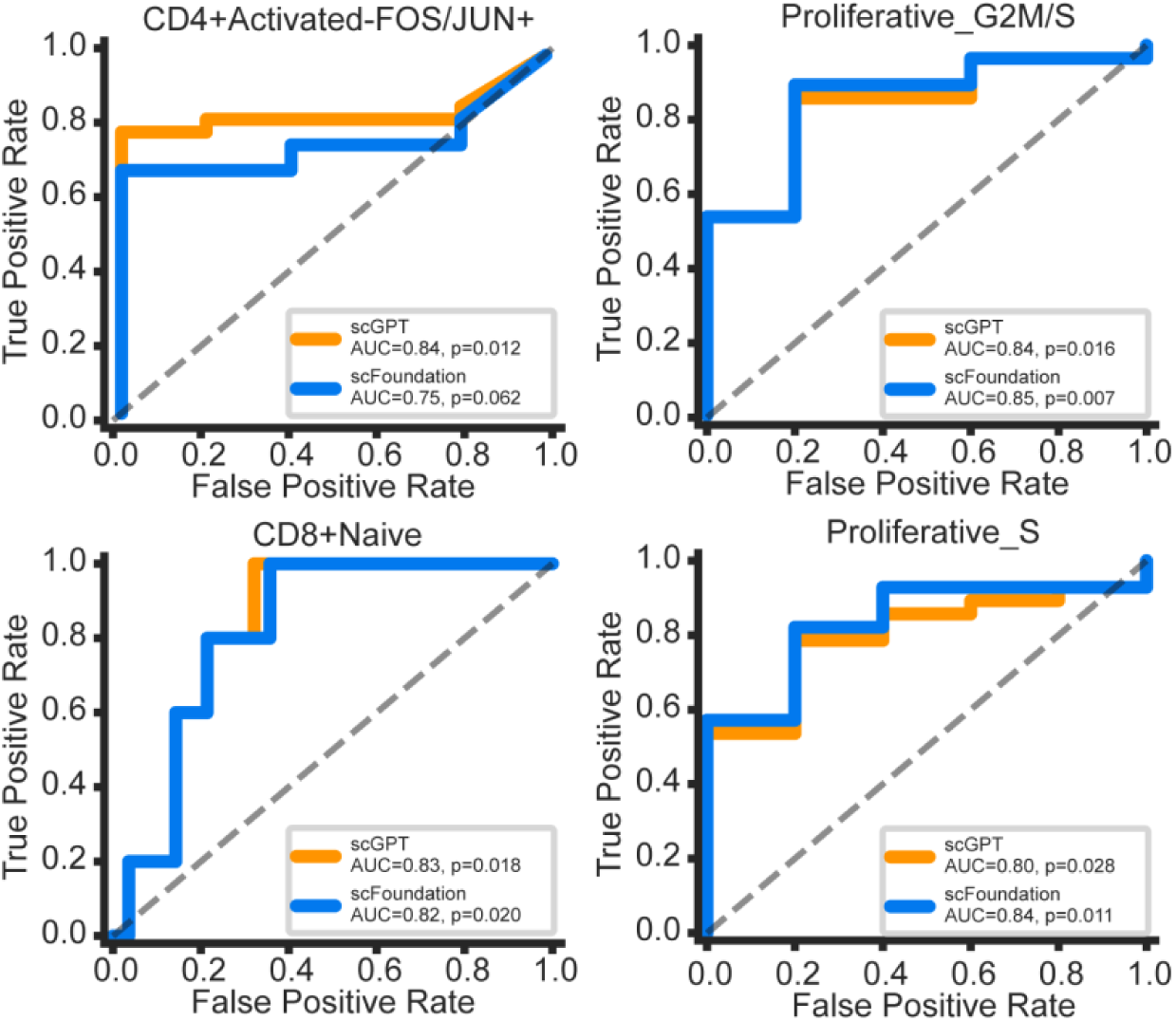
Receiver Operating Curve (ROC) analysis of association between CAR T cell basal states and clinical response. ROC curves for BCA-L vs. BCA1+BCA2 classification using proportions of the four top-performing cell types under basal (unstimulated) condition, for scGPT and scFoundation fine-tuned models.

**Supplementary Fig. 4:**
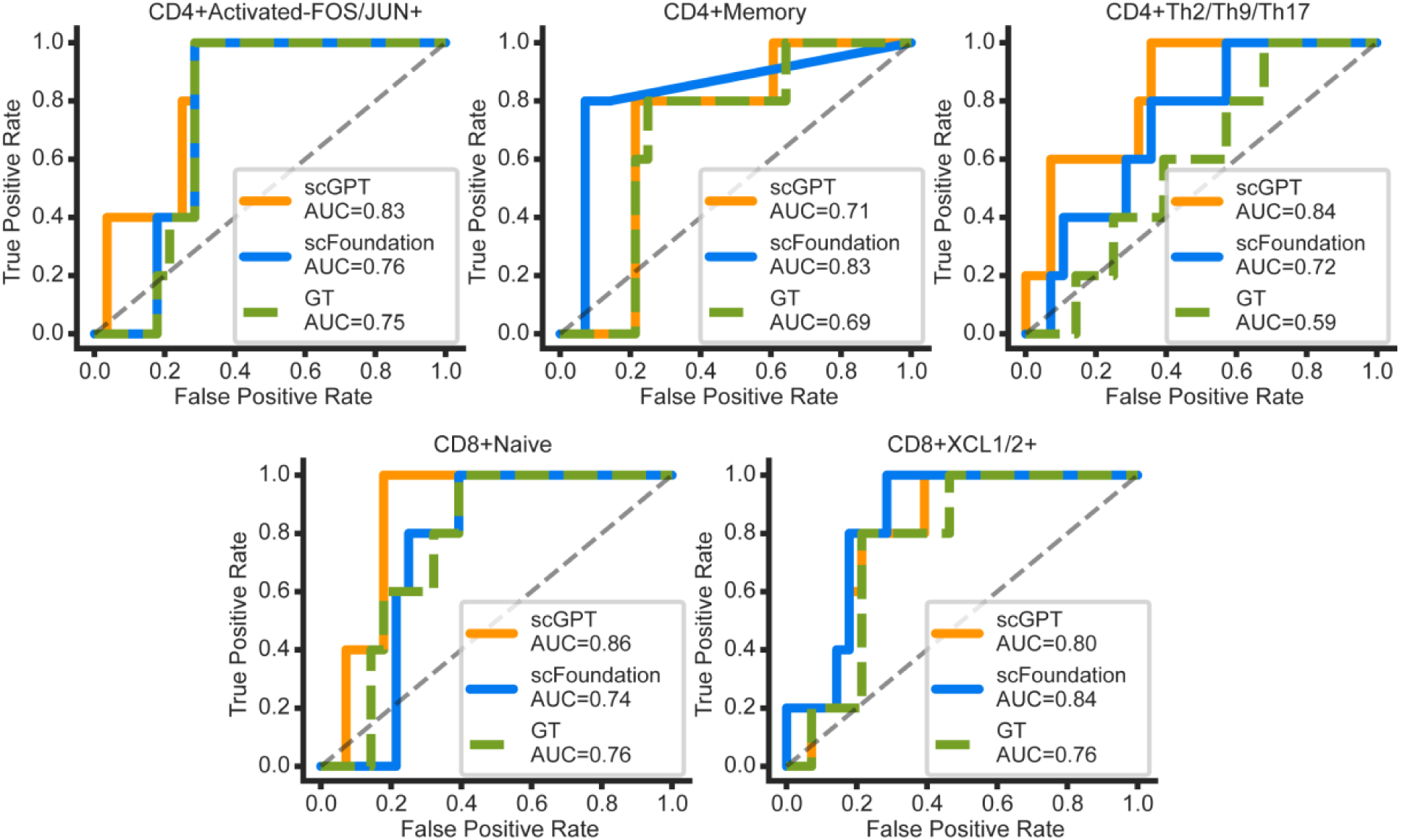
Receiver Operating Curve (ROC) analysis of association between CAR T cell activation states and clinical response. ROC curves for BCA-L vs. BCA1+BCA2 classification under CD19 stimulation condition comparing cell type proportions derived from scGPT fine-tuned, scFoundation fine-tuned, and ground-truth annotations, for the five cell types with highest discriminative performance. AUC values and permutation p-values are shown in each panel.

## References

1 Maude, S. L., et al. Tisagenlecleucel in Children and Young Adults with B-Cell Lymphoblastic Leukemia. New England Journal of Medicine 378, 439–448 (2018). 10.1056/NEJMoa1709866

2 Maude, S. L., et al. Chimeric Antigen Receptor T Cells for Sustained Remissions in Leukemia. New England Journal of Medicine 371, 1507–1517 (2014). 10.1056/NEJMoa1407222

3 Lee, D. W., et al. T cells expressing CD19 chimeric antigen receptors for acute lymphoblastic leukemia in children and young adults: a phase 1 dose-escalation trial. The Lancet 385, 517–528 (2015). 10.1016/S0140-6736(14)61403-3

4 Park, J. H., et al. Long-term follow-up of CD19 CAR therapy in Acute Lymphoblastic Leukemia. New England Journal of Medicine. 378, 449–459 (2018). 10.1056/NEJMoa1709919

5 Schultz, L. M., et al. Disease Burden Affects Outcomes in Pediatric and Young Adult B-Cell Lymphoblastic Leukemia After Commercial Tisagenlecleucel: A Pediatric Real-World Chimeric Antigen Receptor Consortium Report. Journal of Clinical Oncology 40, 945–955 (2022). 10.1200/JCO.20.03585

6 Pulsipher, M. A., et al. Next-Generation Sequencing of Minimal Residual Disease for Predicting Relapse after Tisagenlecleucel in Children and Young Adults with Acute Lymphoblastic Leukemia. Blood Cancer Discovery 3, 66–81 (2021). 10.1158/2643-3230.BCD-21-0095

7 Finney, O. C., et al. CD19 CAR T cell product and disease attributes predict leukemia remission durability. Journal of Clinical Investigation 129, 2123–2132 (2019). 10.1172/JCI125423

8 Bai, Z., et al. Single-cell CAR T atlas reveals type 2 function in 8-year leukemia remission. Nature 634, 702–711 (2024). 10.1038/s41586-024-07762-w

9 Fraietta, J. A., et al. Determinants of response and resistance to CD19 chimeric antigen receptor (CAR) T cell therapy of chronic lymphocytic leukemia. Nature Medicine 24, 563–571 (2018). 10.1038/s41591-018-0010-1

10 Sheih, A., et al. Clonal kinetics and single-cell transcriptional profiling of CAR-T cells in patients undergoing CD19 CAR-T immunotherapy. Nature Communications 11, 219 (2020). 10.1038/s41467-019-13880-1

11 Deng, Q., et al. Characteristics of anti-CD19 CAR T cell infusion products associated with efficacy and toxicity in patients with large B cell lymphomas. Nature Medicine 26, 1878–1887 (2020). 10.1038/s41591-020-1061-7

12 Good, C. R., et al. An NK-like CAR T cell transition in CAR T cell dysfunction. Cell 184, 6081–6100.e26 (2021). 10.1016/j.cell.2021.11.016

13 Cui, H., et al. scGPT: toward building a foundation model for single-cell multi-omics using generative AI. Nature Methods 21, 1470–1480 (2024). 10.1038/s41592-024-02201-0

14 Hao, M., et al. Large-scale foundation model on single-cell transcriptomics. Nature Methods 21, 1481–1491 (2024). 10.1038/s41592-024-02305-7

15 Wen, H., et al. CellPLM: Pre-training of Cell Language Model Beyond Single Cells. In Proceedings of the International Conference on Learning Representations (ICLR) (2024). 10.1101/2023.10.03.560734

16 Rosen, Y., et al. Universal Cell Embeddings: A Foundation Model for Cell Biology. Preprint at bioRxiv (Cold Spring Harbor Laboratory) (2023). 10.1101/2023.11.28.568918

17 Baek, S., Song, K. & Lee, I. Single-cell foundation models: bringing artificial intelligence into cell biology. Experimental & Molecular Medicine. 57, 2169–2181 (2025). 10.1038/s12276-025-01547-5

18 Theodoris, C. V., et al. Transfer learning enables predictions in network biology. Nature 618, 616–624 (2023). 10.1038/s41586-023-06139-9

19 Yang, F., et al. scBERT as a large-scale pretrained deep language model for cell type annotation of single-cell RNA-seq data. Nature Machine Intelligence 4, 852–866 (2022). 10.1038/s42256-022-00534-z

20 Kedzierska, K. Z., Crawford, L., Amini, A. P. & Lu, A. X. Zero-shot evaluation reveals limitations of single-cell foundation models. Genome Biology 26, 101 (2025). 10.1186/s13059-025-03574-x

21 Boiarsky, R., et al. Deeper evaluation of a single-cell foundation model. Nature Machine Intelligence 6, 1443–1446 (2024). 10.1038/s42256-024-00949-w

22 Liu, T., Li, K., Wang, Y., Li, H. & Zhao, H. Evaluating the Utilities of Foundation Models in Single-Cell Data Analysis. Preprint at bioRxiv (2024). 10.1101/2023.09.08.555192

23 Hao, Y., et al. Integrated analysis of multimodal single-cell data. Cell 184, 3573–3587.e29 (2021). 10.1016/j.cell.2021.04.048

24 Li, X., et al. Single-Cell Transcriptomic Analysis Reveals BCMA CAR-T Cell Dynamics in a Patient with Refractory Primary Plasma Cell Leukemia. Molecular Therapy 29, 645–657 (2021). 10.1016/j.ymthe.2020.11.028

25 Salter, A. I. et al. Comparative analysis of TCR and CAR signaling informs CAR designs with superior antigen sensitivity and in vivo function. Science Signaling 14, eabe2606 (2021). 10.1126/scisignal.abe2606

26 Bai, Z., et al. Single-cell multiomics dissection of basal and antigen-specific activation states of CD19-targeted CAR T cells. Journal for Immunotherapy of Cancer 9, e002328 (2021). 10.1136/jitc-2020-002328

27 Böttcher, J. P., et al. NK Cells Stimulate Recruitment of cDC1 into the Tumor Microenvironment Promoting Cancer Immune Control. Cell 172, 1022–1037 (2018). 10.1016/j.cell.2018.01.004

28 Li, Z., et al. Functional Diversification and Dynamics of CAR-T Cells in B-ALL Patients. Cell Reports 42, 113263 (2023). 10.1016/j.celrep.2023.113263

29 Dorner, B. G., et al. Selective Expression of the Chemokine Receptor XCR1 on Cross-Presenting Dendritic Cells Determines Cooperation with CD8⁺ T cells. Immunity 31, 823–833 (2009). 10.1016/j.immuni.2009.08.027

30 Bachem, A., et al. Superior antigen cross-presentation and XCR1 expression define human CD11c⁺CD141⁺ cells as homologues of mouse CD8⁺ dendritic cells. Journal of Experimental Medicine 207, 1273–1281 (2010). 10.1084/jem.20100348

31 Bai, Z., et al. Single-cell antigen-specific landscape of CAR T infusion product identifies determinants of CD19-positive relapse in patients with ALL. Sience Advances 8, eabj2820 (2022). 10.1126/sciadv.abj2820

32 Lopez, R., Regier, J., Cole, M. B., Jordan, M. I. & Yosef, N. Deep generative modeling for single-cell transcriptomics. Nature Methods 15, 1053–1058 (2018). 10.1038/s41592-018-0229-2

33 Lotfollahi, M., et al. Mapping single-cell data to reference atlases by transfer learning. Nature Biotechnology 40, 121–130 (2022). 10.1038/s41587-021-01001-7

34 Cui, H. (subercui). The number of parameters in scGPT. GitHub issue #125, bowang-lab/scGPT repository (2024). https://github.com/bowang-lab/scGPT/issues/125

35 DiCiccio, T. J. & Efron, B. Bootstrap confidence intervals. Statistical Science 11, 189–228 (1996). 10.1214/ss/1032280214

36 Benjamini, Y. & Hochberg, Y. Controlling the False Discovery Rate: A Practical and Powerful Approach to Multiple Testing. Journal of the Royal Statistical Society: Series B (Methodological) 57, 289–300 (1995). 10.1111/j.2517-6161.1995.tb02031.x

